# Privacy Preserving Epigenetic PaceMaker Stronger Privacy and Improved Efficiency

**DOI:** 10.1101/2024.02.15.580590

**Authors:** Meir Goldenberg, Loay Mualem, Amit Shahar, Sagi Snir, Adi Akavia

## Abstract

DNA methylation data plays a crucial role in estimating chronological age in mammals, offering real-time insights into an individual’s aging process. The Epigenetic Pacemaker (EPM) model allows inference of the biological age as deviations from the population trend. Given the sensitivity of this data, it is essential to safeguard both inputs and outputs of the EPM model. In a recent study by Goldenberg et al., a privacy-preserving approach for EPM computation was introduced, utilizing Fully Homomorphic Encryption (FHE). However, their method had limitations, including having high communication complexity and being impractical for large datasets Our work presents a new privacy preserving protocol for EPM computation, analytically improving both privacy and complexity. Notably, we employ a single server for the secure computation phase while ensuring privacy even in the event of server corruption (compared to requiring two non-colluding servers in Goldenberg et al.). Using techniques from symbolic algebra and number theory, the new protocol eliminates the need for communication during secure computation, significantly improves asymptotic runtime and and offers better compatibility to parallel computing for further time complexity reduction. We have implemented our protocol, demonstrating its ability to produce results similar to the standard (insecure) EPM model with substantial performance improvement compared to Goldenberg et al. These findings hold promise for enhancing data security in medical applications where personal privacy is paramount. The generality of both the new approach and the EPM, suggests that this protocol may be useful to other uses employing similar expectation maximization techniques.

## 1 Introduction

Genomic data is protected by privacy regulations, such as the European General Data Protection Regulation (GDPR) [27], California Customer Privacy Act (CCPA) [26], and Genetic Information Privacy Act (GIPA) [11] that may limit the ability of companies and researchers to collect large cohorts of genomic data as required for training machine learning models of high predictive power. A promising approach for overcoming such limitations is to execute *privacy preserving genome analysis*, i.e., to utilize cryptographic techniques such as fully homomorphic encryption (FHE) [29,15] that enable processing data in encrypted form, so that even the entity processing it is never exposed to the underlying data in cleartext. Prior work on privacy preserving genome analysis using homomorphic encryption focused primarily on Genome Wide Association (GWAS) [24,32,8,7,12] –i.e., statistically associating innate genome variability in single nucleotide polymorphism (SNPs) with a risk for a disease or a particular trait– as well as on privacy preserving classification of DNA and RNA sequences of tumor tissues [20,10] and viral strains [41,2] respectively. Recently, privacy preserving epigenetics –i.e., the study of how behavior and environmental factors lead to genome changes that in turn affect the phenotype– was considered in [3,16], analysing gene expression data [3] and DNA methylation data [16].

Elaborating on the latter, DNA methylation are chemical changes in the genome that are linked to numerous developmental, physiologic, and pathologic processes including malignancy, infections, and aging. A breakthrough in this area was achieved by the *epigenetic clock* of Horvath [21] that predicted human chronological age based on observing several hundreds of methylation sites along the genome. Seeking to link between these methylation processes and aging, Snir et al. [36] suggested a flexible, probabilistic framework adopted from the evolutionary realm [37,40,38], under which a maximum likelihood solution is sought, called the Epigenetic PaceMaker (EPM). The output of the algorithm is a point in the likelihood surface under which the probability of the entire system is optimised while preserving basic model constraints. The flexibility of the framework enabled explaining intrinsic key features in aging, both in human and animals [30,28]. In [35] an Expectation Maximization algorithm was proposed to search the likelihood surface via several iterations consisting of two steps, each increasing the likelihood function. Subsequently, using algebraic identities, along with the special structure of the EPM framework, closed form solutions to both steps were found, yielding asymptotic reduction in time and space complexities [33]. One of the outputs of the EPM framework is the *epigenetic state* (also referred to as the “biological age”, to be contrasted with the chronological one). The epigenetic state can provide significant information on the health status of an individual, which can be utilized, for example, in estimating the efficacy of pharmaceutical interventions or in predicting life expectancy. Accurate estimations and predictions may require however a larger dataset than available for a single entity such as a hospital or a research lab, making it desirable to combine datasets from several entities, albeit, without compromising privacy. To address the privacy issue a first step was made in Goldenberg et al. [16], proposing a privacy preserving protocol for computing the EPM model using FHE as a central tool. Their protocol required two servers throughout the entire computation, resulting in two main drawbacks: first, their protocol is insecure if an adversary corrupts both servers; second, their protocol suffers from a high communication and computational toll, making it impractical to real data volumes.

In this work we propose a new privacy-preserving protocol for the EPM, improving over the prior work [16] in the following aspects:

– **Improved complexity: no communication**. In [16], secure computing entails multiple rounds of communication between the two servers: one communication round for each expectation-maximization iteration in computing the EPM, transmitting *O*(*nm*) ciphertexts in each iteration (for *n* the number of methylation sites, and *m* the number of individuals in the dataset). In contrast, our secure computing entails *no communication* between the two servers, regardless of the number of expectation-maximization iterations securely computed (cf. Section 3.1).^1^
– **Stronger privacy: hiding both input and output**. In [16], the produced epigenetic ages are revealed to all parties. In contrast, we offer an enhanced security version of our protocol where we *protect the privacy of both input and output* (cf. Section 3.2). That is, rather than revealing all epigenetic ages to all parties, we reveal each epigenetic age only to the entity who provided the data for the corresponding individual. This strengthening requires only a single round of interaction between the servers, transmitting a single ciphertext in each direction. Moreover, if we slightly relax the output hiding requirement as to allow exposing the common denominator of the produced epigenetic ages, then this privacy enhancement incurs no interaction.
– **Stronger privacy: relying on a more plausible assumption**. Both ours and [16]’s security guarantee relies on an assumption that the two servers are non-colluding, i.e., an adversary can corrupt one server or the other, but not both. In [16] both servers are required to be online and actively participate throughout the entire protocol, including during the computing phase which entails the bulk of the computation; this often makes a non-collusion assumption challenging to realize. In contrast, in our protocol only one server executes the secure computing phase, while the other server’s role is essentially minimized to be the role of a key management service (KMS), i.e., key generation and decryption in the pre- and post-processing phases, respectively, whereas it can be offline and idle throughout the secure computing phase. Being offline throughout the bulk of the computation makes it considerably easier to safeguard this server, thus increasing the plausibility of the non-collusion assumption.^2^

We implemented our protocol and show that it offers a substantial performance speed-up over the implementation of [16]: securely computing the EPM from 716 methylation sites in 3 hours (cf. 24 sites in 3 hours in [16]). These results can serve as a pilot for data security measures integrated in vast medical applications where personal privacy is imperative.

In terms of techniques, we focus on offering a new algorithm for computing the EPM model, which avoids the complexity bottlenecks of the underlying FHE (such as computing matrix inverse and division). Concretely, our algorithm substitutes matrix inverse computations with closed-form algebraic formulas that were computed analytically building on [33]; and substitutes division computations by representing rational numbers as pairs of integers – their numerator and denominator. We note that the above interventions lead to high magnitude numbers throughout the computation, which we reduce to a manageable scale using the Chinese Remainder Theorem.

## 2 Preliminaries and Definitions

### 2.1 The Epigenetic Pacemaker (EPM): Model and Algorithm

We summarize the EPM model and optimization problem [36]. Let *s*_1_, …, *s*_*n*_ be *methylation sites* in the genome that exist in every individual and undergo methylation during life at characteristic *rate r*_*i*_. Each site *s*_*i*_ is initiated at birth with an *initial level* of methylation, denoted 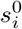. Both 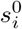 and *r*_*i*_ are universal - common to all individuals. Actual rate at a specific individual may however vary. However the *EPM property* mandates that site rates change proportionally throughout lifetime over all sites of the same individual. That is, at any point in time, if a change in rate occurred in site *i* of individual *j*, then the rates in *all* sites *i*^*′*^ in *j*, are simultaneously changed and by the same factor, so that the ratio between any two rates *r*_*i*_ and 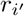 is maintained at all times. While methylation rates might even be negative (demethylation) it is these changes in the rates that are correlated with the *aging rate*, providing a good estimate on the *epigenetic age (e-age)* of an individual (as opposed to its chronological age)[36]. We denote by *t*_*j*_ the weighted average e-age of individual *j*, accounting for the rate changes an individual has undergone through life.

The algorithmic task under the EPM model is to find the maximum likelihood values of 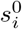, *r*_*i*_, and *t*_*j*_, when given the observed methylation levels in *n* genome sites as measured in *m* individuals. The input is denoted by (*ŝ*_*i,j*_)_*j*∈[*m*],*i*∈[*n*]_ where *ŝ*_*i,j*_ denotes the methylation measured in individual *j* at site *i* (where [*x*] = 1, 2, 3, …, *x*).

An algorithm for the EPM optimization problem was presented by Snir et al. [36], who show that their algorithm provably converge to a local optima of the maximum likelihood function. Subsequently, [33] presented an experimental evaluation validating the concrete good efficiency of this algorithm. This algorithm is the starting point of our solution.

To describe the algorithm we first organize the observed variables *ŝ*_*ij*_ as well as the unknown variables *t*_*j*_, *r*_*i*_ and 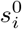 as follows. Let *X* be a *mn* × 2*n* matrix whose *k*th row is all zero except for the value *t*_*j*_ in the *i*th entry of its first half and 1 in the *i*th entry of its second half. Let *β* be a column vector whose first *n* entries are *r*_1_, …, *r*_*n*_ and the last *n* entries are 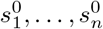. Let *y* be the column vector whose *im* + *j* entry contains *s*_*i,j*_. See Figure 5 in the Supplementary Material.

The algorithm of [33] consists of several iterations, where in each iteration the algorithm alternates between two main components: a *site step* and a *time step*. In the site step values for *t*_*j*_’s are fixed to be the values obtained from the previous iteration (on the first iteration, they are initialized to random values), and the algorithm solves the linear regression problem system specified by *X, y* (where *X* is with the said values for *t*_*j*_’s) to obtain values from *r*_*i*_ and 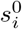(i.e., for *β*). In the time step, *r*_*i*_ and 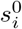 are fixed to the values obtained in the site step (of the current iteration), and the individual’s times are set to their maximum likelihood values, which as proved by [33], is given by the following closed form rational function:

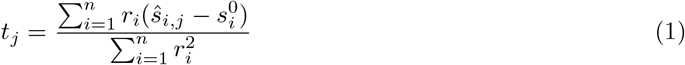

Furthermore, [33] proves that at every such step an increase in the likelihood is guaranteed, and so, a local optimum is eventually reached. The iterations can proceeds until the improvement in the Residual Sum of Square (RSS) falls below a threshold *δ* given as a parameter to the algorithm.

Our goal is to compute the epigenetic age in a *privacy preserving* fashion. Therefore, we must not reveal even the number of iterations required for convergence, because this could potentially reveal significant information on the input (see [3] for discussion of such attacks). We therefore slightly modify the algorithm from [33], in specifying the number of iterations in advance, by a user-defined parameter denoted iter. This algorithm is summarized in Figure 6 in the Supplementary Material.

### 2.2 Privacy Preserving EPM Computation over Federated Data: Problem Definition

#### Privacy preserving EPM

*settings and goal*. In a privacy preserving EPM computation over federated data addresses, we consider a setting in which *m* individuals, called *Data Owners* and denoted by DO_1_, …, DO_*m*_, each data owner DO_*j*_ holds observed methylation levels *ŝ*_1,*j*_, … *ŝ*_*n,j*_ in *n* sites *s*_1_, …, *s*_*n*_.^3^ The data owners wish to compute the epigenetic age estimator specified by the EPM algorithm on their joint data, but without revealing information on their individual data. The data owners encrypt the data and outsource the computation to a server called the Machine Learning Engine (MLE) who executes the computation, whereas the complexity of the data owners is proportional only to the size of their individual input (in encrypted form). The parties also have access to a Crypto Service Provider (CSP), that can generate key pairs and decrypt for authorized values (alternatively, provide decryption keys to authorized parties). The goal is to compute the same epigenetic age estimation as outputted by the EPM algorithm when executed on the union of the individual data, but without exposing any information on the raw data (beyond what can be inferred from the designated output and leakage profile). This is summarized in the *EPM functionality* depicted in Figure 1.

**Fig. 1:**
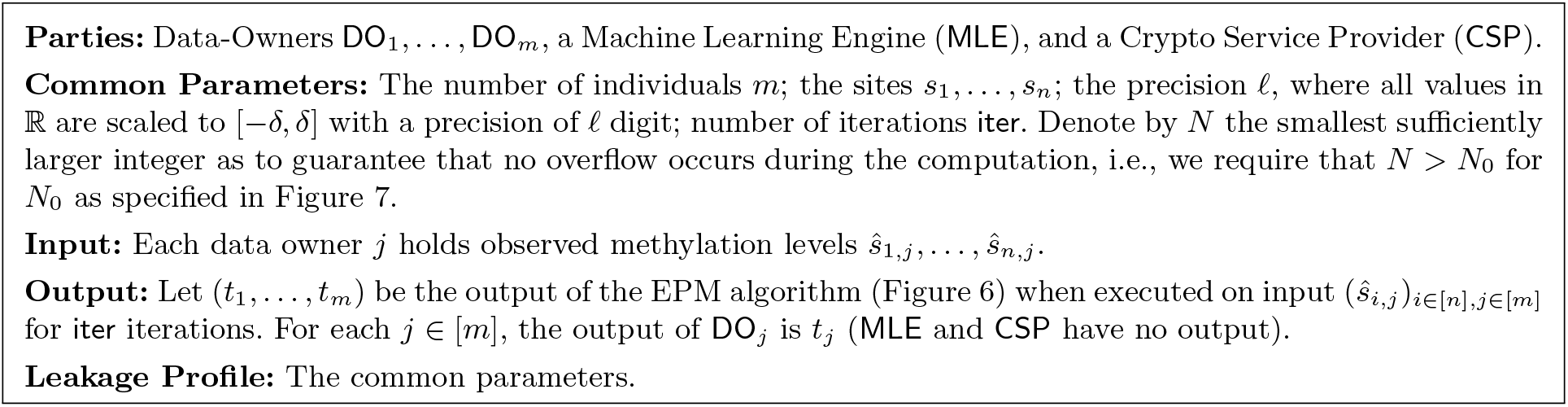
EPM Functionality

#### Threat model and security requirement

To achieve the above goal, the parties engage in an interactive protocol in which parties can repeatedly send messages to each other and execute local computations on their input and received messages. We analyze security in the two-server model, as in [16]; that is, the security requirement is to guarantee correctness and privacy against all passive computationally-bounded adversaries who may corrupt any subset of the data owners and at most one out of the CSP and MLE. Being passive means that parties controlled by the adversary follow the protocol specification, albeit they may collude to infer as much information as possible from their view of the interaction. Being computationally-bounded means that all parties are restricted to performing probabilistic polynomial time computations.

To capture this formally we first specify some standard terminology. Let Π be a protocol for computing EPM; denote by *x*_1_, …, *x*_*m*_ the inputs of DO_1_, …, DO_*m*_; and *λ* the security parameter. The output in an execution of Π on these inputs and security parameter is a random variable denoted by output^Π^ (*x*_1_, …, *x*_*m*_) (where the probability here is over the randomness of all participating parties, including the servers). The output of the EPM functionality (cf. Figure 1) on these inputs is a random variable denoted by EPM(*x*_1_, …, *x*_*m*_) (where the probability is over the randomness of the EPM algorithm (cf. Figure 6)), and we denote by EPM(*x*_1_, …, *x*_*m*_)_*j*_ its *j*th coordinate which is output of DO_*j*_. The *correctness* requirement is that with overwhelming probability the output of the protocol is identical to the output of EPM (see Definition 1, Correctness). The *view* of any party *P* ∈ {DO_1_, …, DO_*m*_, MLE, CSP} during an execution of Π on inputs *x*_1_, …, *x*_*m*_ of DO_1_, …, DO_*m*_ respectively and security parameter *λ* is the random variable consisting of the *input* and *randomness* of *P* and the *messages P received* from the other parties during the execution of Π, and denoted by 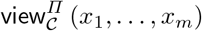. The *privacy* requirement is that for any set *𝒞* of corrupt parties that includes at most one of the two servers, the view of *𝒞* is computationally indistinguishable from a random variable that can be efficiently computed when given only the input and output of corrupt parties and the leakage ℒ (as specified in Figure 1). This captures the property that participating in the protocol does not equip the corrupt parties with any further knowledge. See Definition 1, Privacy.

##### Definition 1 (Securely realizing EPM)

*We say that a protocol* Π securely realizes the EPM functionality *(cf. Figure 1) with leakage* − *against passive computationally-bounded adversaries in the two-server model if the following holds:*

1. ***Correctness:*** *There exists a negligible function negl* (*λ*) : ℕ → ℕ *such that for all inputs* 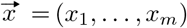 *and security parameter λ*,

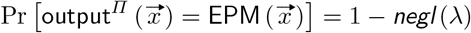

*(where the probability is over the randomness of the parties in the protocol and over the randomness of the EPM algorithm)*.
2. ***Privacy:*** *For every set of corrupt parties* 𝒞 ⊂ {*DO*_1_, …, *DO*_*m*_, *MLE, CSP*} *consisting of any number of data owners and at most one of MLE and CSP, there exists a computationally bounded simulator* Sim *such that for every input* 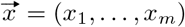,

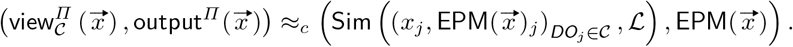

*where* ≈_*c*_ *denotes that these two random variables are* computationally indistinguishability.^4^

### 2.3 Fully Homomorphic Encryption

We use fully homomorphic encryption (FHE) [29,15] as a key tool in our protocol. FHE supports encrypting messages and processing the resulting ciphertexts –without knowledge of the underlying messages– to obtain ciphertext for the results of computations on these underlying messages. For example, given two ciphertexts *c*_1_ and *c*_2_ encrypting messages *m*_1_ and *m*_2_ it is possible to produce ciphertexts *c*_Add_ and *c*_Mult_ so that decrypting these ciphertexts produces the messages *m*_1_ + *m*_2_ and *m*_1_ *· m*_2_ respectively. The ring in which the arithmetic operations are computed is called the *plaintext space*. We employ an FHE that supports, for any integer *N* ≥ 2, the plaintext space ℤ_*N*_, i.e., the ring integers modulo *N*, and where *N* is provided as input during key generation; we refer to this *N* as the *plaintext modulus*.

More formally, a FHE scheme is a 4-tuple of probabilistic polynomial time (aka, ppt) algorithms *E* = (KeyGen, Enc, Dec, Eval) with the following syntax: KeyGen takes as input a security parameter *λ* and an integer *N* ≥ 2 and outputs a key pair (*pk, sk*) ← KeyGen(1^*λ*^, *N*). Enc takes as input a public key *pk* and a message msg ∈ ℤ_*N*_, and outputs a ciphertext ctxt ← Enc(*pk*, msg). Dec takes as input a secret *sk* and a ciphertext ctxt, and outputs a plaintext message msg^*′*^ ← Dec(*sk*, ctxt). The correctness requirement is that for every (*pk, sk*) in the range of KeyGen, with overwhelming probability: Dec(*sk*, Enc(*pk*, msg)) = msg. Eval takes as input a public key *pk*, a function *C* accepting *k* inputs and producing *l* outputs for some *k, l*, and a vector of *k* ciphertexts 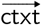, and outputs a vector of *l* ciphertexts 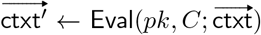. The *homomorphism* requirement is that for all (*pk, sk*) in the range of KeyGen, with overwhelming probability, Dec(*sk*, Eval(*pk, C*; Enc(*pk*, msg_1_), …, Enc(*pk*, msg_*k*_))) = *C*(msg_1_, …, msg_*k*_)) *Compactness* says that the ciphertext size is independent of the class of supported homomorphic computations. The *semantic security* requirement is that for every *λ, N* and msg ∈ ℤ_*N*_, the joint distribution of *pk* (generates by (*pk, sk*) ← KeyGen(1^*λ*^, 1^*N*^)) and ctxt ← Enc (*pk*, msg) is computationally indistinguishable from the joint distribution of *pk* and ctxt_0_ ← Enc (*pk*, 0).

## 3 Our Privacy Preserving EPM Protocol

In this section, we describe our privacy-preserving protocol for computing the EPM over federated data. For simplicity of the presentation, in Section 3.1 we first describe a protocol for a simplified functionality in which the privacy of the input is protected whereas the output (the e-ages) are revealed to all parties. Moreover, these e-ages (which are rational numbers) are represented in the output by a pair of integers – their numerator and denominator. Subsequently, in Section 3.2, we describe the expansion of this protocol to conceal both the input and the output, i.e., to securely realize the EPM functionality (Figure 1).

The protocol is executed by the following parties. Data owners DO_1_, …, DO_*m*_, where each DO_*j*_ has as input the observed methylation levels *ŝ*_1,*j*_, … *ŝ*_*n,j*_.^5^ A Machine Learning Engine Server (MLE) which performs the secure protocol computation. A Crypto Service Provider (CSP) who provides the public encryption and evaluation keys to the data owners and MLE, respectively, and decrypts the output.^6^ All parties know the common parameters, which consist of the number of individuals *m*, the number of methylation sites *n*, the precision 𝓁, an upper bound *τ* on the e-ages, and the employed homomorphic encryption scheme *ε* = (KeyGen, Enc, Dec, Eval). The FHE *ε* supports homomorphic computation with plaintext arithmetic over ℤ_*N*_ (i.e., where the message space consist of integers and the arithmetic operations –addition and multiplication– are computed modulo *N*). The plainatext modulus *N* is of size *O*(iter *·* (𝓁 + log_2_(*mnτ*)), is efficiently computed from the common parameter (see the formal details in the Supplementary Material, Figure 7).

The security and complexity guarantees of our protocol are specified in the following theorems. We use the *O*_*λ*,log *N*_ () notation to hide complexity terms that are polynomial in *λ* and *N* and are determined solely by the complexity of the underlying encryption scheme *E* while being irrespective to our protocol; and denote by slots the number of data items that can be packed in each ciphertext. The proof is provided in the Supplementary Material.

### Theorem 1 (Security)

*Our protocol securely realizes the EPM functionality (Figure 1) against any passive computationally-bounded adversary controlling any number of the data owners and at most one of MLE and CSP*.

### Theorem 2 (Complexity)

*The complexity of our protocol is as follows*.

– *Each DO*_*j*_*’s runtime is dominated by the time to encrypt n methylation values, i*.*e*., *O*_*λ*,log *N*_ (*n*).
– *The CSP’s runtime is dominated by the time to decrypt the m (masked) output values, i*.*e*., *O*_*λ*,log *N*_ (*m*).
– *The MLE’s runtime is dominated by the time to homomorphically compute an arithmetic circuit of size iter · O* (*m*^2^ + *nm*) *and multiplicative-depth* 3(*iter* − 1).
– *The communication volume is dominated by the size of the encrypted input, i*.*e*., *O*_*λ*,log *N*_ (*mn*).
– *The secure computing phase involves 1-round of interaction between the MLE and CSP (and is non-interactive, if revealing the common denominator of the output values is permitted)*.

### 3.1 Our Simplified Protocol: Hiding the Input

Our input hiding EPM protocol is presented next (see also Figures 7-8 in the Supplementary Material). The protocol is composed of three phases –pre-processing, secure computing, and output post-processing– as detailed next.

*Phase I: Pre-processing*. The pre-processing phase consists of key setup and data upload. Key setup is executed by the CSP who generates the required FHE key pairs and publishes the public keys. Data upload is executed by the DO_*s*_ who format their methylation data, encrypt it, and pass the ciphertexts to the MLE for the secure computation. Data upload can occur over a period of time where methylation levels are gradually uploaded in encrypted form to cloud storage that will be made accessible to the MLE for secure computation when needed. Formatting includes rounding the data to 𝓁 decimal digits (for the 𝓁 specified in the common parameters), scaling the rounded values to integers (e.g., if 𝓁 = 2 and the methylation values are in (−1, 1) then scaling is via multiplying each value by 100), and utilizing the Chinese Remainder Theorem (CRT), following [4], for homomorphically processing plaintext values too large to be directly supported by existing FHE libraries (see Remark 1).

*Remark 1 (CRT representation)*. To address the issue of large (plaintext) values arising in our protocol we rely on the Chinese Remainder Theorem (CRT), following [4], as follows. We set the plaintext modulus *N* to be a product of smallish co-prime numbers *N* = *p*_1_ *· p*_2_ … *· p*_*k*_ (e.g., 60 prime numbers of size 30-bits each, in our empirical evaluation), and rely on the isomorphism 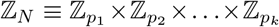 to replace the computation modulo *N* with *k* computations modulo *p*_1_, *p*_2_, …, *p*_*k*_ respectively. To support this CRT approach, the CSP generates *k* key pairs (*pk*_1_, *sk*_1_) ← KeyGen(1^*λ*^, 1^*p*^1), …, (*pk*_*k*_, *sk*_*k*_) ← KeyGen(1^*λ*^, 1^*p*^*k*) and publishes *pk* := (*pk*_1_, …, *pk*_*k*_); the data owners encrypt each methylation value *ŝ*_*ij*_ under all keys, producing vectors of ciphertext Enc(*pk, ŝ*_*ij*_) := (Enc(*pk*_1_, *ŝ*_*ij*_), …, Enc(*pk*_*k*_, *ŝ*_*ij*_)) that are passed to the MLE; and the MLE executes the homomorphic evaluat#i o n« with# respect to «all keys (in parallel), i.e., homomorphic evaluation of a function *C* on cipher 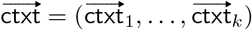 is by computing 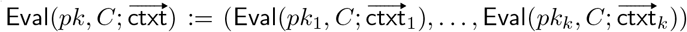. For the sake of readability in the following (including in Figures 7-8), and simply write *pk*, Enc(*pk, ·*) and Eval(*pk, ·*; *·*).

*Phase II: Secure computing*. In the secure computing phase the MLE employs homomorphic evaluation to produce a ciphertext encrypting the estimated e-ages, and publishes those ciphertexts. The homomorphic computation produces the same e-ages as in the EPM algorithm (cf. Figure 6), but includes important modifications that bypass complexity bottlenecks associated with homomorphic computation. To provide details, first recall that the EPM algorithm executes several iterations, each consisting of a *site step*, in which matrix inversion is computed to solve a linear regression problem recovering latent parameters (called the rate 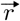 and the methylation at birth 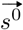), and a *time step*, in which division is computed to update the e-ages 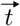 using these latent parameters. Both matrix inversion and division suffer from poor efficiency when computed homomorphically, and our solution avoids both, as follows. First, to avoid homomorphic matrix inverse we directly compute the solution to the regression problem using a closed-form algebraic solution from [33, Lemma 3]. This formula however entails division, a complexity bottleneck that we avoid as discussed next. Second, to avoid homomorphic division we *represent rational numbers by the pair of integers: their numerator and denominator*, where we homomorphically update these (integral) values throughout the computation. That is, we represent rational numbers 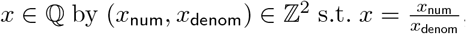, and dividing *x* by an integer *Λ* is done by homomorphically updating its denominator to be*· Λ*^−1^. These mathematical reformulations of the calculations, in both the site step and the time step phases, result in a formulation that exclusively involves addition, subtraction, and multiplication operations – which are efficient to compute homomorphically with FHE. We note that keeping track of the growing numerator and denominator, rather than computing division, leads to high magnitude numbers throughout the computation, which we reduce to a manageable scale using the Chinese Remainder Theorem (as discussed above).

*Phase III: Output post-processing*. In the output post-processing phase, the CSP (or any authorized party that can fetch *sk* from CSP) decrypts and publishes the e-ages numerators and denominator (if the denominator is zero, output ⊥). Optionally: CSP performs division over the reals to obtain the cleartext e-ages in the standard representation of rational numbers.

### 3.2 Our Full Protocol: Hiding Both Input and Output

For simplicity of the presentation the protocol described above reveals the entire vector of e-ages to all parties. It may be desired however to increase privacy so that each e-age (*t*_num,*j*_, *t*_denom_) is exposed in cleartext only to the corresponding data owner DO_*j*_, while being hidden from all other parties. Moreover, it may be desired to hide also the denominator *t*_denom_ and reveal to each DO_*j*_ only the ratio 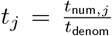 (where division is over the reals). This can be achieved via minor modifications. In what follows, we explain how to extend the protocol to hide also the output values 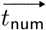 and *t*_denom_, except the designated output receiver. We first explain how to hide 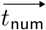, and then how to also hide *t*_denom_.

*Hiding the numerators of the e-ages*. Revealing each (*t*_num,*j*_, *t*_denom_) only to DO_*j*_ (and to no other party) is done as follows. In the data upload phase, each data owner DO_*j*_ samples a uniformly random mask mask_*j*_ ∈ ℤ_*N*_, encrypts it under *pk*, and sends the ciphertext to the MLE along with the encrypted methylation values. In the secure computing phase, before publishing the e-ages, the MLE homomorphically masks each e-ages numerators *t*_num,*j*_ by computing ***t***^***′***^_num,*j*_ ← Eval(*pk*, Add; ***t***_num,*j*_, **mask**_*j*_) (where the function Add, given two integers, outputs their sum modulo *N*), and sends to the CSP these masked values 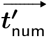 along with ***t***_denom_. In the output post-processing phase, the CSP decrypts and sends to each DO_*j*_ his masked output 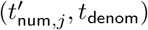 in cleartext. Each DO_*j*_ then unmasks by computing 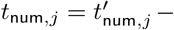mask_j_ mod *N* (in cleartext), and outputs 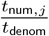 where division is over the reals (output ⊥ if *t*_denom_ = 0).

*Hiding also the denominator*. The denominator *t*_denom_ is the same for all DO_*j*_’s, and so producing the output in the form of a pair of numerator and denominator cannot hide the denominator. When desired to hide also *t*_denom_, it can be done by adding the following further modifications to the above. In the secure computing phase, MLE does not publish 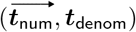 but rather does the following. MLE samples uni-formly random mask 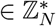 and homomorphically masks *t*_denom_ by ***t***^***′***^_denom_ ← Eval(*pk*, Mult; ***t***_denom_, **mask**) (where Mult is the function that, given two integers, outputs their product modulo *N*), and sends the masked encrypted value ***t***^***′***^_denom_ to the CSP. The CSP then decrypts, computes (in cleartext) the inverse of 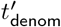 in ℤ_*N*_ (except for outputting ⊥, if 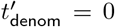), denoted 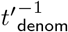, encrypts and sends the re-sulting ciphertext 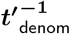 to MLE. In response, the MLE first homomorphically unmasks this inverse by computing 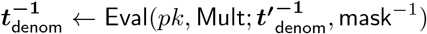 (where mask^−1^ is the inverse of mask in ℤ_*N*_, which MLE computes in cleartext); second, homomorphically computes 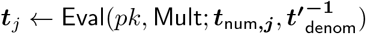, for each *j* ∈ [*m*]; third, homomorphically masks the resulting values with the mask_*j*_ received from the data owners by computing ***t***^***′***^_*j*_ ← Eval(*pk*, Add; ***t***_*j*_, **mask**_***j***_), and sends the resulting masked ciphertexts 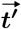 to the CSP. In the output post-processing phase, the CSP decrypts, and, for each *j* ∈ [*m*], sends 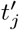 to DO_*j*_. Subsequently, for each *j*, DO_*j*_ unmasks to produce 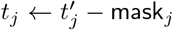, and then computes rational reconstruction [39,14] to produce the rational number that is equal to computing 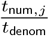 over the reals.

## 4 System Description and Empirical Evaluation

To assess the precision and efficiency of the secure protocol we proposed, we put it into practice as a proof of concept. An overview of our protocol’s procedural flow is depicted in Fig 2.

**Fig. 2:**
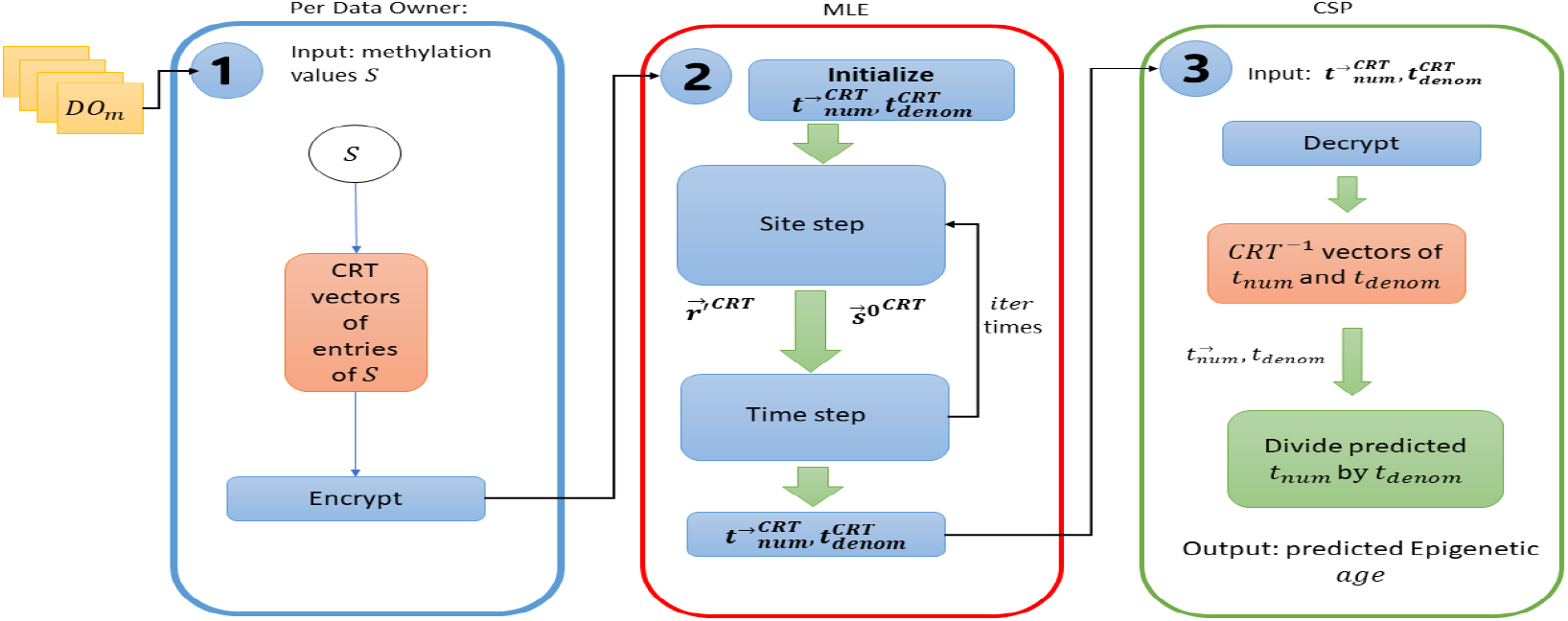
System flow: (1) Each Data Owner employs the Chinese Remainder Theorem (CRT) on her methylation values (S) and encrypts the resulting CRT representation (aka the CRT vector). Denote by 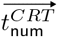 and 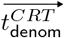 the CRT vectors of 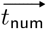 and *t*_denom_. (2) MLE initializes 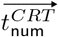 and 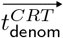 executes iter iterations of the site and time steps. (3) The final output is the predicted epigenetic ages.

*Library versions*. We implemented our proof of concept version of the protocol in python version 3.10.12 using Pyfhel version 3.4.1 [22] as the FHE library, sympy version 1.12 [25] for CRT and numpy version 1.24.3 [19] for matrix storage and operations.

*Data set*. For our real data experiment, we used the GSE74193 describing DNA methylation patterns in the frontal cortex during development. The methylation profile was generated by [23] and used in [34] to demonstrate logarithmic growth of human epigenetic aging along the lifespan.

*Data preparation*. The encrypted methylation values and ages provided by the data owners are represented as floating-point numbers with an abundance of digits in the fractional part. However, our selected Fully Homomorphic Encryption (FHE) scheme supports only integer data type, and so we convert the data (methyaltion and age values) to integers via scaling and rounding. To minimize the loss of accuracy, we conducted tests using various rounding values. Our experimentation revealed that rounding the numbers to 2 digits in the fractional part would result in an accuracy loss of approximately 0.5% for the majority of the outcomes.

For our initial feature selection phase, we utilized Pearson Correlation, a standard technique in machine learning. This process involved removing sites from the training data that displayed low correlation with the target ages. We found that an 80% correlation coefficient led to the selection of 716 sites, which is considered a suitable number for our proof of concept. In this context, precision, determined by the number of digits (𝓁), was set to 2, rounding the decimal fraction of the numbers to two digits. The values were scaled within the range [−*δ, δ*), with *δ* having a maximum value of 10000. This constraint arises from the consideration that all ages in our dataset fall within the range of 0 to 99 years. The dataset comprises a total of *m* = 472 individuals.

*Encryption parameters*. FHE has several parameters that require configuration to match various use cases. Throughout the implementation phase, we conducted a thorough assessment of various parameter configurations and encryption schemes, ultimately arriving at the following set of parameters, which align well with the volume of mathematical operations and data dimensions used in our implementation and testing.

We use the Brakerski/Fan-Vercauteren (BFV) scheme [9], [13] with a 16k polynomial modulus degree, 30 bit plaintext modulus prime numbers and a security key size of 128 bits.

Furthermore, our implementation leverages the packing feature, an integral component of FHE encryption. This feature enables the encryption of multiple cleartext values within a single encrypted context object, granting us the capability to perform mathematical operations on all values in parallel. This provides a significant increase in calculation efficiency in cases of vector multiplication and addition operations as well as calculation of several individual values in parallel as required in both the site and time steps. Pyfhel aligns the packed array size to the polynomial modulus degree [1], encompassing 16k individual cells in a 2 × 8*k* matrix format. As the method for accessing the second row of cells in the matrix did not conform well with our implementation requirements, we utilized only the first 8k of cells. Required changes have since been added to Pyfhel and can be a point for improvement of our implementation in the future.

*Multiplicative depth*. In order to guarantee the security of FHE, each ciphertext must contain a certain level of noise. Mathematical operations, particularly multiplication, contribute significantly to noise accumulation. If the noise surpasses a predefined threshold, it results in an overflow, rendering the ciphertext indecipherable. The threshold is determined by encryption context parameters, with the polynomial modulus degree and plaintext modulus value being the most influential. Our protocol’s analysis determined the maximum number of consecutive multiplications, termed *multiplicative depth*, performed on a ciphertext within a single iteration. Our analysis revealed a multiplicative depth of 3 for the calculation of ***t***_num_ which increases with each iteration of the protocol. This prompted exploration for noise mitigation solutions. While increasing the polynomial modulus degree is an option, it adversely affects performance and memory requirements. An alternative is *bootstrapping*, a noise reduction operation. Although time-consuming, if applied selectively, it minimally impacts the overall protocol execution time. Our analysis suggested that applying bootstrapping to ***t***_num_ and ***t***_denom_ at the end of the time step, should support any amount of iterations. Unfortunately, our chosen FHE library lacked bootstrapping support, leading us to simulate it through a decrypt-re-encrypt process with a trusted third party. This approach is for testing purposes only and falls short of meeting our strict security requirements. Future authentic bootstrapping support is essential for securely implementing the protocol. Meanwhile, bootstrapping tests in C++ using HElib [18] yielded a 159-second execution time for a single 16k polynomial modulus degree which must be considered as part of the overall protocol execution time.

*Handling large values*. The necessity to avoid division operations on the encrypted ciphertexts results in the accumulation of very large integer values which grow with each protocol iteration. To mitigate this challenge, we adopt the Chinese Remainder Theorem (CRT) for representing each input value. CRT allows us to decompose integers into the residue classes modulo pairwise coprime moduli, facilitating more efficient computations. Applying CRT to these integer values effectively addresses the representation size limitation practically imposed by the encryption parameters. However, the implementation of the CRT approach posed challenges, particularly when dealing with negative values.

Basic operations such as subtraction or multiplication can alter the sign of a value. Despite this, due to the modulo operations inherent in our system, our value image range is confined to [0, *N* − 1], where *N* is the product of all primes used. Formally, for a given set of primes *𝒫*, there exists two distinct integers, *x* and *y*, such that −*N < y <* 0 *< x < N*, yet *x* mod *𝒫* = *y* mod *𝒫*. To tackle this sign issue, we follow [6,31], and use an additional bit. This extra bit can be employed as a sign bit, ensuring that *x* mod *𝒫* and *y* mod *𝒫* possess unique representations.

*Parallel computation*. As previously mentioned, our approach entails multiple algorithm runs, each with a different prime, to calculate the final result via CRT. We conducted a performance evaluation by executing 3 iterations of the protocol with a single prime number, revealing a total runtime of 45 minutes. In order to mitigate the substantial increase in runtime that would arise from running with multiple prime numbers in serial, we implemented parallel multi-process computation. In this setup, each process is responsible for the computation associated with a single prime number.

*Result figures*. our program was run on an AWS cloud server featuring 96 Intel Xeon 8275CL CPUs, 192GB of RAM and Ubunutu Sever 22.04 OS. We used 60 prime numbers for the computation, thereby concurrently executing 60 parallel processes each running three iterations of the protocol, resulting in an observed execution time of 3 hours and 7 minutes. To ascertain an accurate depiction of the time required for the dataset under consideration, we must incorporate the estimated time of the bootstrapping operations. As we expected additional overhead due to parallel execution, we tested 60 parallel bootstrap operations and observed a total runtime of 282 seconds. As there are four bootstrap operations in three iterations of the protocol, we added 4 × 282 seconds to the total expected execution time which amounts to: 3 hours, 25 minutes and 48 seconds.

We proceeded to perform a comparative analysis by assessing the ages computed by our secure protocol in relation to those obtained through our implementation of the clear-text algorithm. We measured the difference in percentage between the e-ages calculated by both implementations. The error per individual *i* is calculated by: 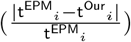 where t^EPM^ and t^Our^ represent the ages calculated by the cleartext algorithm and our secure protocol accordingly. The results are depicted in Figure 3. As can be evidenced, for 446 out of 472 individuals, the difference between the secure and cleartext protocol is 0.5% or less. For an additional 16 individuals, the difference is between 0.5% to 1% and for the remaining 10 individuals, the difference is between 1% to 5%.

**Fig. 3:**
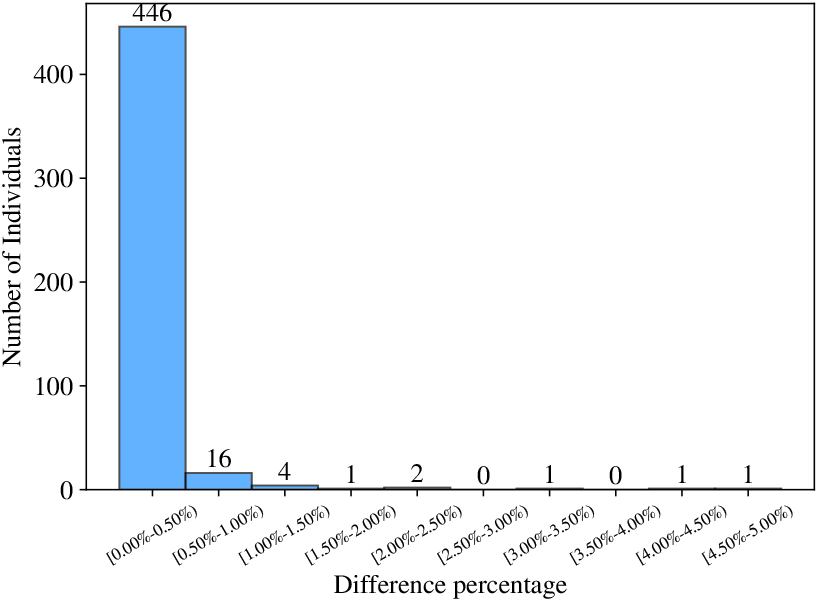
Epigenetic age differences percentage in our protocol vs. EPM over cleartext. The *x* axis depicts error rate ranges; the *y* axis depicts the number of samples (individuals) fitting the error rate.

Furthermore, we measure how execution time scales in the number of sites, by measuring the runtime of our protocol also on 24 sites (as in [16]) and 1514 sites, all with 472 individuals. The results are depicted in Figure 4b.

**Fig. 4:**
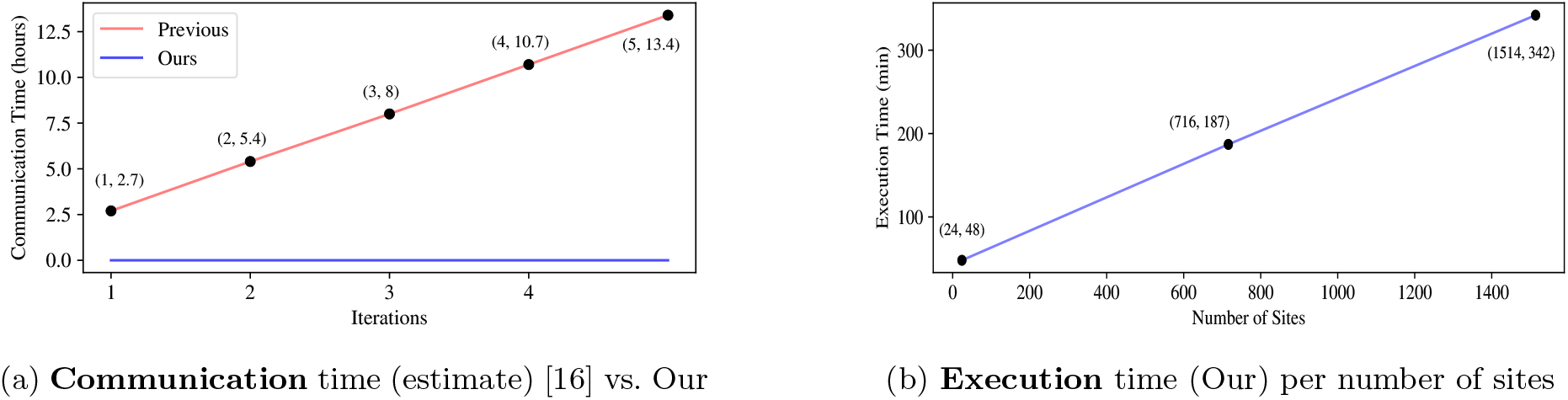
Computing phase complexity: communication and computing time.

**Fig. 5:**
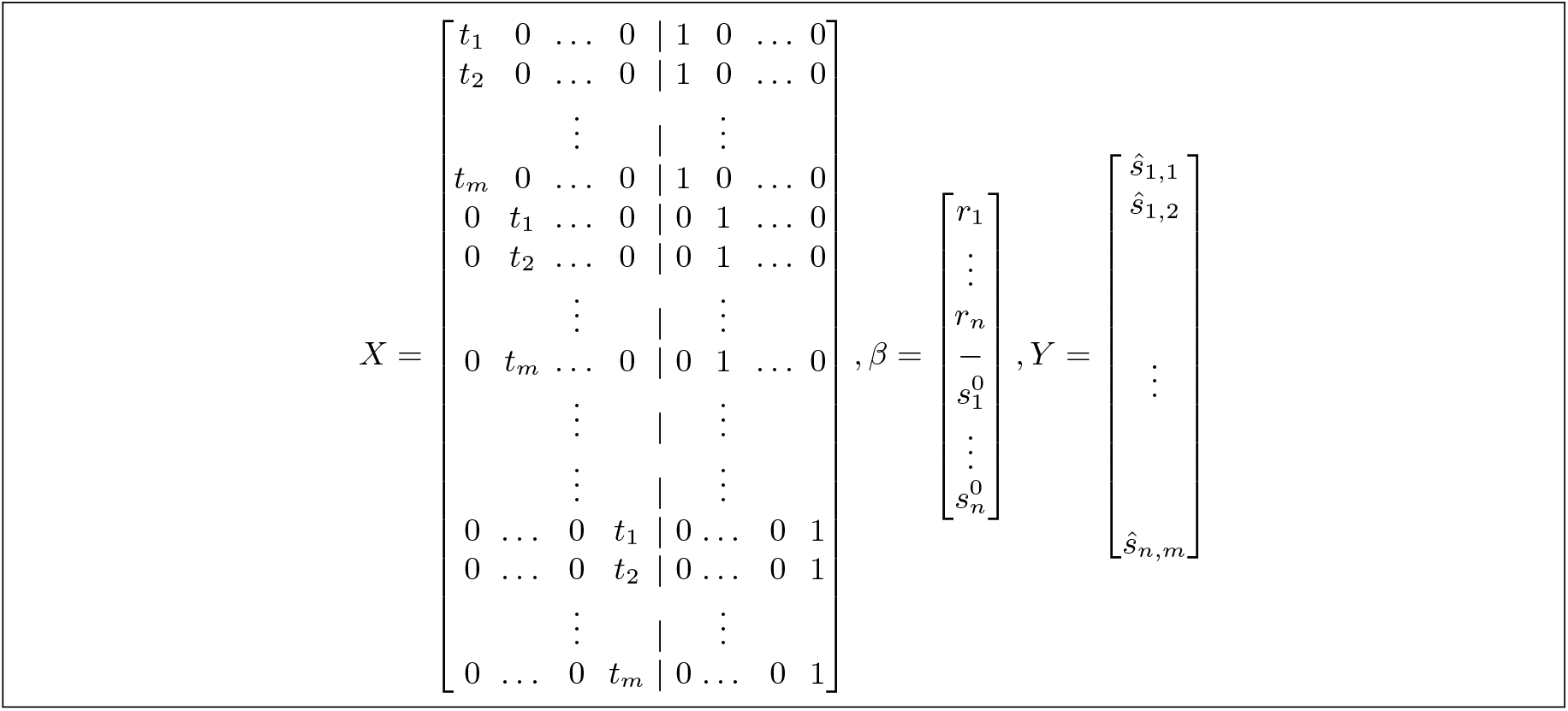
*X, β*, and *y*

**Fig. 6:**
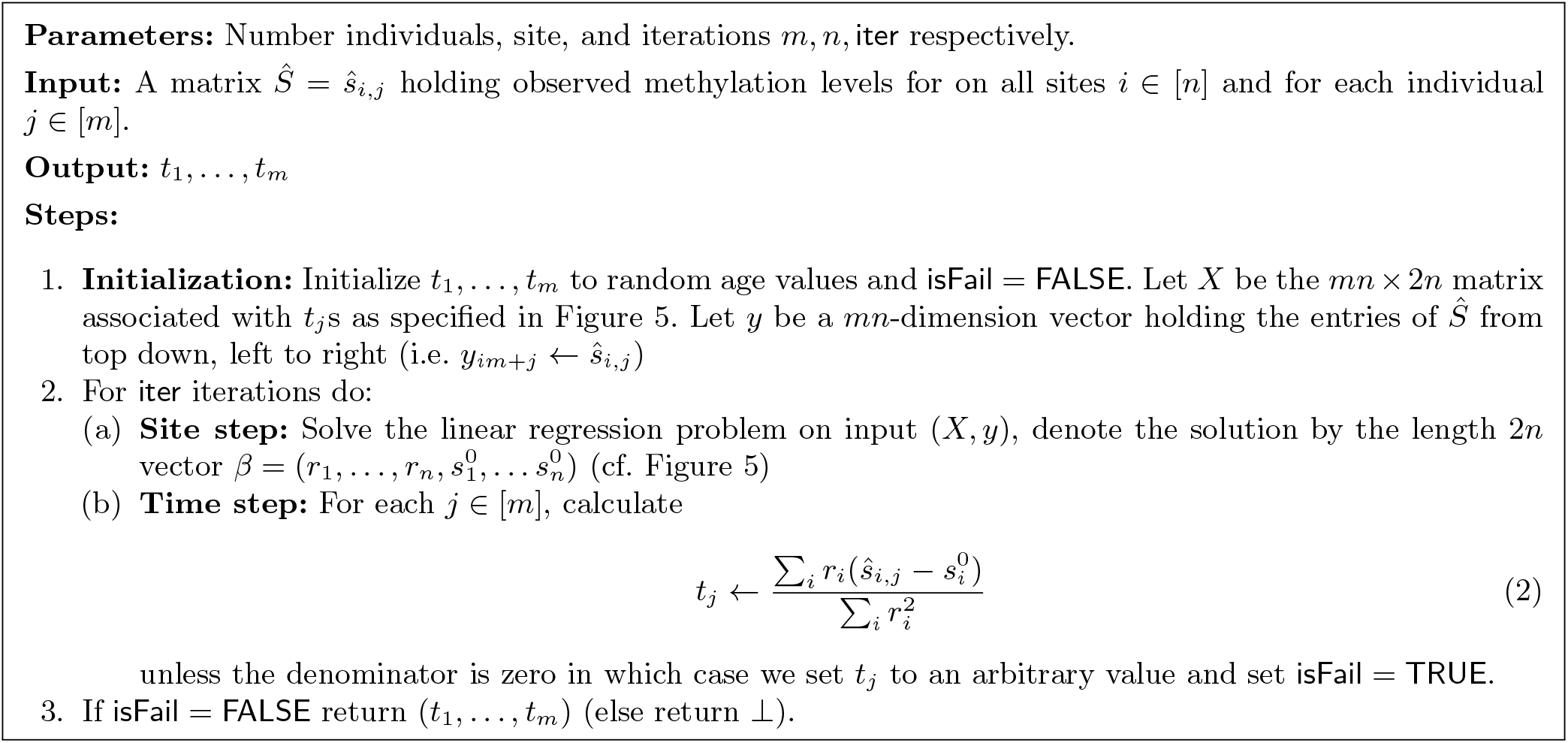
EPM Algorithm on cleartext data

**Fig. 7:**
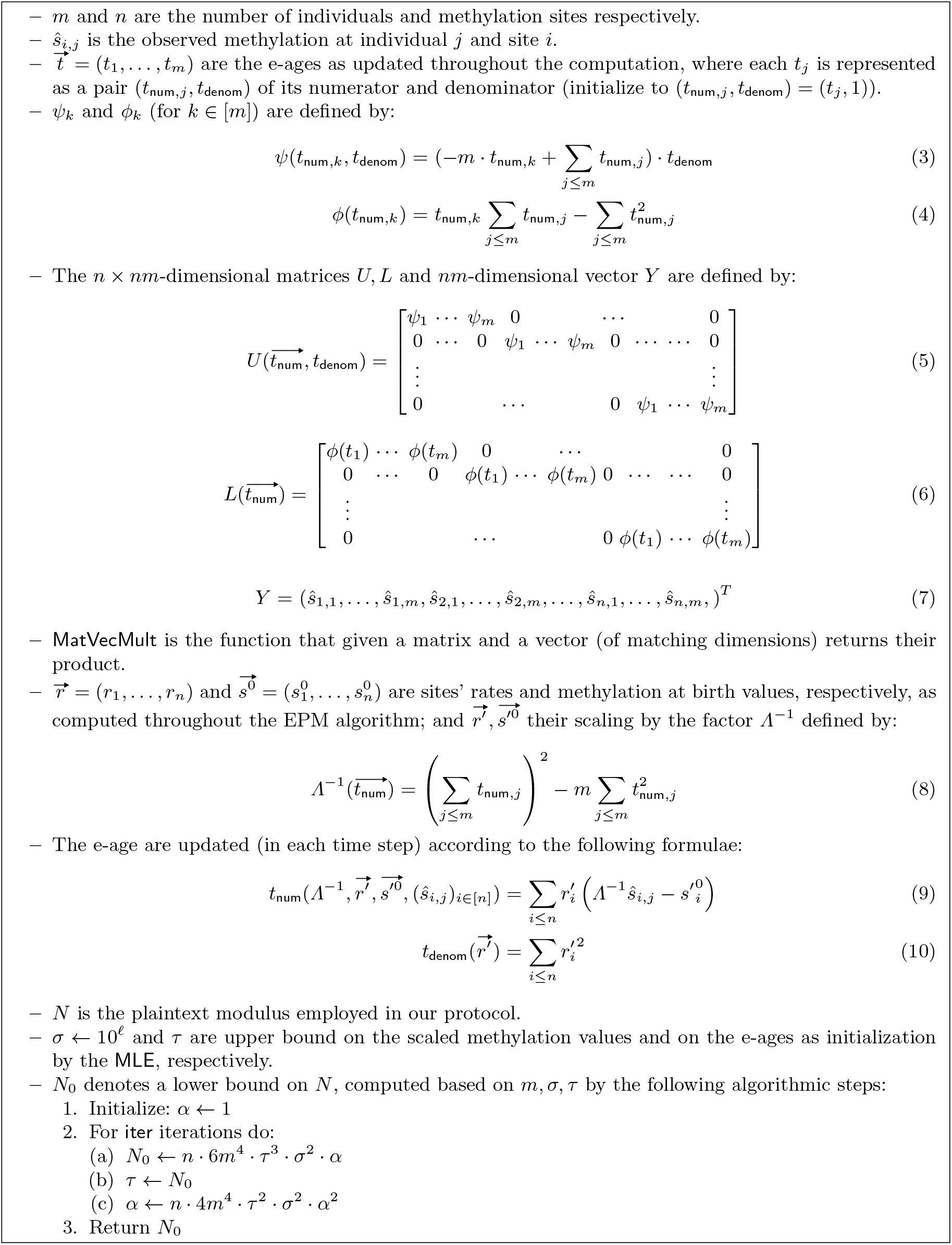
Notations summary: inputs, variables, and functions to be homomorphically evaluated in our protocol.

**Fig. 8:**
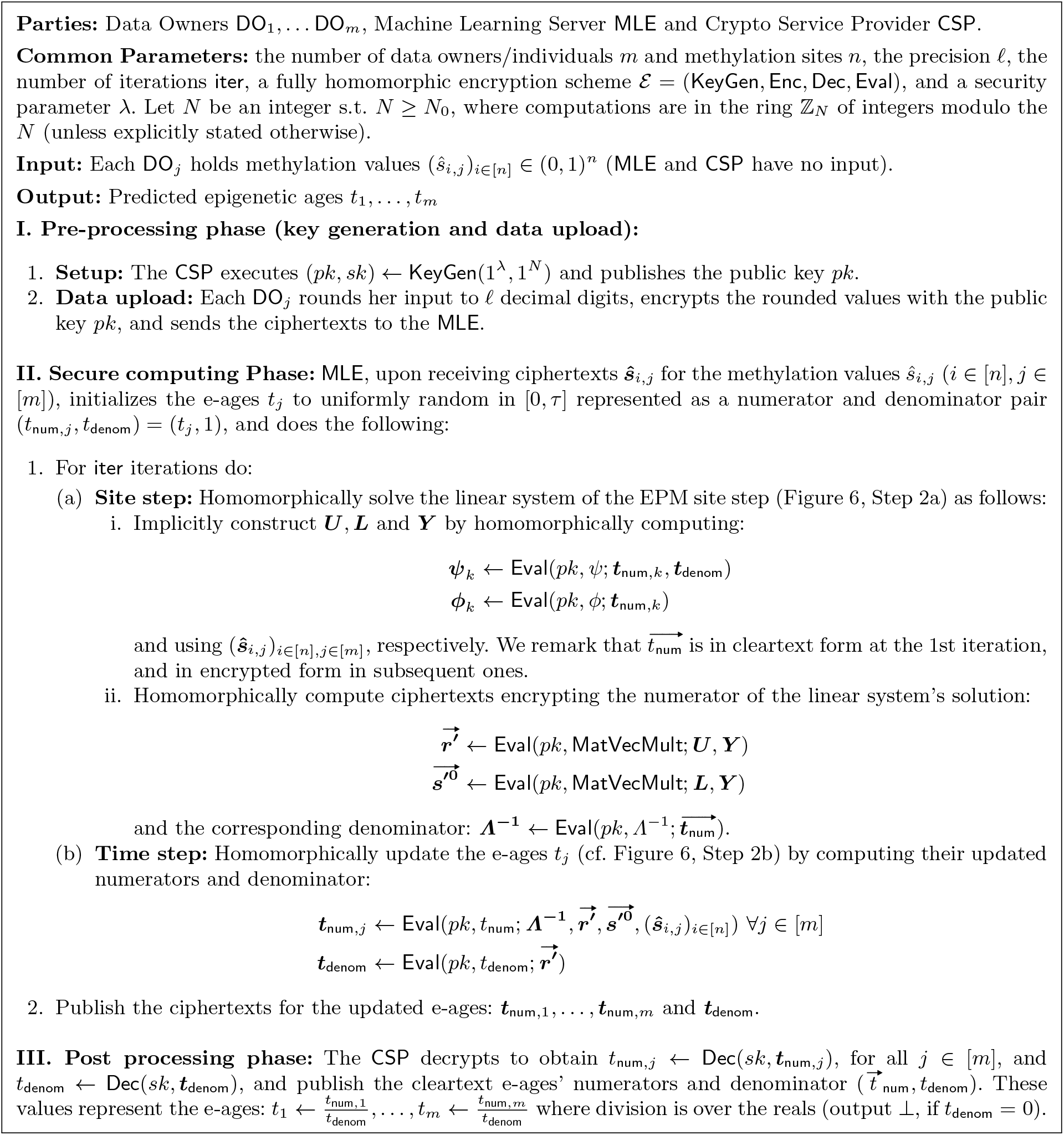
Our Secure EPM Protocol. See Figure 7 for the definition of the functions *Λ*^−1^, *ψ, ϕ*, MatVecMult, *t*_num_,*t*_denom_, matrices *U, L*, vector *Y*, and « the bounds *τ* and *N*_0_. Ciphertexts are denoted in **Boldface**, i.e., 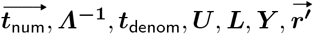 and 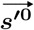 decrypt to 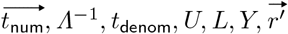 and 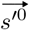 respectively. We note that *Λ*^−1^ denotes both a function and its output 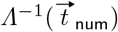.

In addition, we estimate the communication time between the two servers required for executing the protocol of [16]: in each iteration their communication entails the MLE sending an encrypted 2*n* × 2*n* matrix and receiving from the CSP an encrypted 2*n*-dimensional vector, i.e., transmitting a total of iter *·* (4*n*^2^ + 2*n*) encrypted values. As their protocol was not fully implemented, we must complement some implementation details to estimate their their performance. First, we suppose their implementation employs FHE supporting packing SLOTS=16k cleartext values in each ciphertext, to have equal footing with the packing parameter in our implementation. We then calculated the size in bytes of a ciphertext with 16k slots, which resulted in a value of 2MB. Second, we suppose they utilize packing as recommended in the work [5] they rely on, implying that they transmit 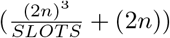 ciphertexts in each iteration (cf. [5, Section 5.2.1, last paragraph]). Third, we suppose their communication is over a Wide Area Network (WAN) of a typical speed (150Mbps); the focus on WAN follows by supposing they would mount their two servers on separate cloud service provides (e.g., AWS vs. Google Cloud) to increase the plausibility of their non-collusion assumption. Supposing all the above, the expected communication time of [16], when executed on the same number of sites, individuals and iterations as in our protocol (i.e., *n* = 716, *m* = 472 and iter = 3), is 8 hours. Further results are depicted in figure 4a.

The github repository of our implementation can be found under: https://github.com/ASEC-lab/EPM-code

## 5 Conclusions

We presented a new privacy preserving protocol for computing the EPM model over federated data. Our protocol offers better asymptotic communication complexity and stronger privacy than the prior work [16] as well as faster concrete efficiency (see Table 1). The uniqueness of the direction pursued here stems from the fact that privacy in the context of epigenetics, as opposed to genome wide association, was rarely studied before, only a single work handling gene expression data [3] preceded our direction of epigenetic aging as pursued here. Moreover, the processing done here is not over sequences, DNA or protein, and hence borrows tools from other areas from the latter. Additionally, the specific conditional expectation maximization structure employed here, by alternating between two phases, in which at each phases simple mathematical operations in the form of closed form solutions developed from well structured, sparse, matrices, may be general enough, leading to other applications, either in life science or even beyond that.

**Table 1:**
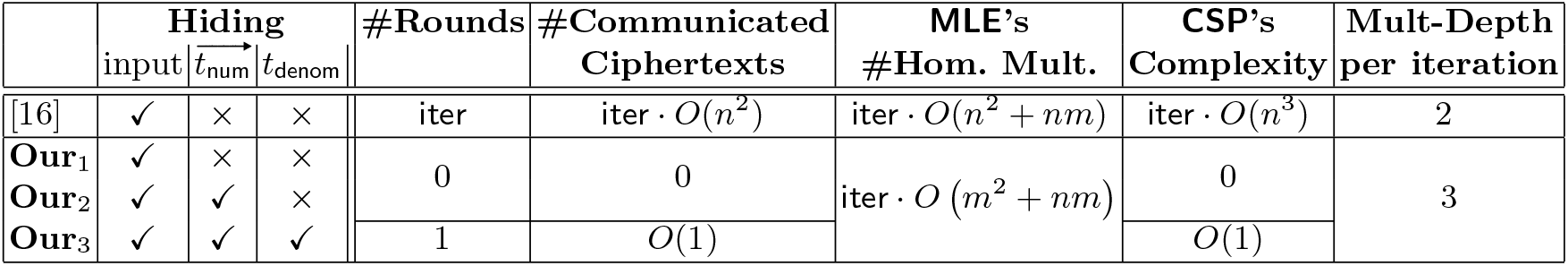
Complexity of the secure computing phase in [16] vs. Ours (three variants). (Here *m, n*, iter are the number of individuals, methylation sites and iterations; and #, Hom. and Mult. are a shorthand for number of, Homomorphic and Multiplication or Multiplicative, respectively.)

## A. Supplementary Material

We provide here formal details that were omitted from the paper due to space constraints. The covered topics are as follows:

– Summary of the EPM Algorithm and Our Privacy Preserving EPM Protocol
– Proof of Theorem 1, Correctness
– Proof of Theorem 1, Privacy
– Proof of Theorem 2, Complexity
– Proof of maximum plaintext modulus value 7
– Encryption and Numeric Representation

### A.1 Summary of the EPM Algorithm and Our Privacy Preserving EPM Protocol

The EPM algorithm is summarized in Figures 5-6; and our privacy preserving protocol for computing EPM over federated data is summarized in Figures 7-8.

### A.2 Proof of Theorem 1, Correctness

*Proof (of Theorem 1, Correctness)*. We hereby sketch the main idea to prove that the ciphertext produced by the MLE upon the termination of the secure computing phase (hencefoth, the secure algorithm) decrypt to the same value as produced by executing the algorithm of Snir [33] (cf. Figure 6) on cleartext values (hencefoth, the cleartext algorithm).

We note that all calculations in the cleartext algorithm are performed over the real numbers while the secure algorithm performs calculations modulo *N* (after proper rounding and scaling of the input), where we assume that *N* is sufficiently large so that no value exceeds *N/*2. The correctness proof is w.r.t to the cleartext algorithm when executed on the values with the same scaling and rounding.

By the correctness of the FHE scheme (cf. Section 2.3), the computations performed homomorphically over encrypted data decrypt to the same value as if applying those computations on the underlying cleartext values. Therefore, it suffices to prove that the latter has the same outcome as the cleartext algorithm. This is not immediate, because the homomorphic computations entail modifications that we introduced to the cleartextg algorithm to avoid complexity bottlenecks. Yet, as we argue below, indeed this is the case.

Snir [33, Lemmas 2 and 3] describes two matrices [*U* ]_*k,l*_ and [*L*]_*k,l*_ of structure similar to our *U, L* (see 5 and 6) except that their non zero diagonal elements are defined to be:

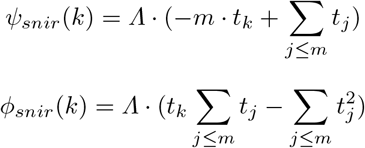

where Λ is defined as:

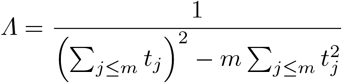

This is similar to our *ψ, ϕ* (see 3,4), except that we avoid the multiplication by *Λ* (because it implies division).

Snir defines the output of the site step to be the product of these matrices [*U* ]_*k,l*_ and [*L*]_*k,l*_ by the vector *Y* as specified in 7. We defined the output analogously, albeit with our matrices *U, L*, leading to a different outcome in our site step than in Snir’s. Denote Snir’s outcome by 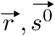 and our outcome by 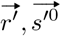, then the following holds:

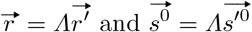

Although our output of the the site step differ from Snir’s, we compensate by modifying the time step accordingly, to have the combination of our site plus time step produce the exact same outcome as the combination of Snir’s site plus time step. Details on our time step follow.

Snir’s time step formula [33, Lemmas 4] calculates the predicted epigenetic age *t*_*snir,j*_ for individual *j* as follows:

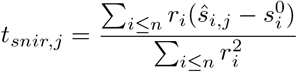

Using the identities 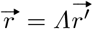 and 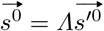 we see that *t*_*snir,j*_ is equal to our *t*_*j*_ that was defined to be:

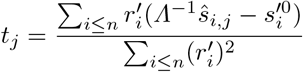

We securely calculate the numerator and denominator per individual *j* denoted by *t*_num,*j*_ and *t*_denom,*j*_ and pass them to the site step for the next iteration. Note that, at this point, our *t*_*j*_ and Snir’s *t*_*snir,j*_ are indeed equal, except that ours are represented by their pair of numerator and denominator. In the final iteration the MLE passes to CSP the ciphertexts for *t*_num,*j*_ and *t*_denom_ who decrypts and divides to compute the output values *t*_*j*_, which –as we have showed– is exactly the same value as Snir’s.

Now we explain how to proceed to the site step of the next iteration. The updated value of *Λ* for the next iteration can be defined using 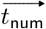 and 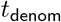 as follows:

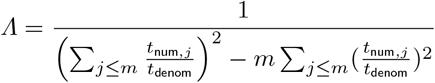

which, following mathematical simplifications, is equivalent to:

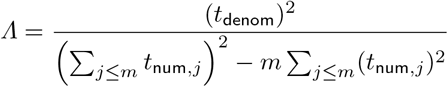

It is noted that in contrast to the initial definition of *Λ* above, the numerator is now (*t*_denom_)^2^ (rather than 1) which needs to be considered during the next iteration. Subsequently, we repeat the site step using the newly computed numerators and denominator for the age values calculated by the time step, multiplied by (*t*_denom_)^2^. Consequently, The updated values of *ϕ* and *ψ* computed in our protocol are equivalent to the following:

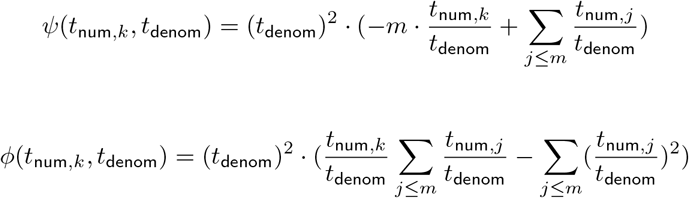

After some mathematical simplifications:

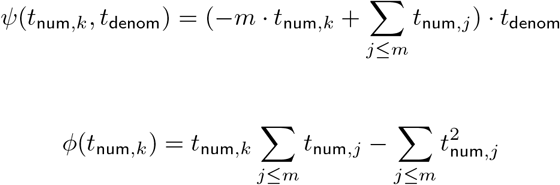

We note that these are identical to the values of *ϕ* and *ψ* in 4, 3. Henceforth, we can proceed to the time step calculation following the same method described above and perform additional iterations if required.

### A.3 Proof of Theorem 1, Privacy

We first prove that the protocol in Figures 7-8 securely realizes the output revealing EPM functionality (Figure 9). We then extend the analysis to show that the extended protocol (cf. Section 3.2 securely realizes the EPM functionality (Figure 1).

**Fig. 9:**
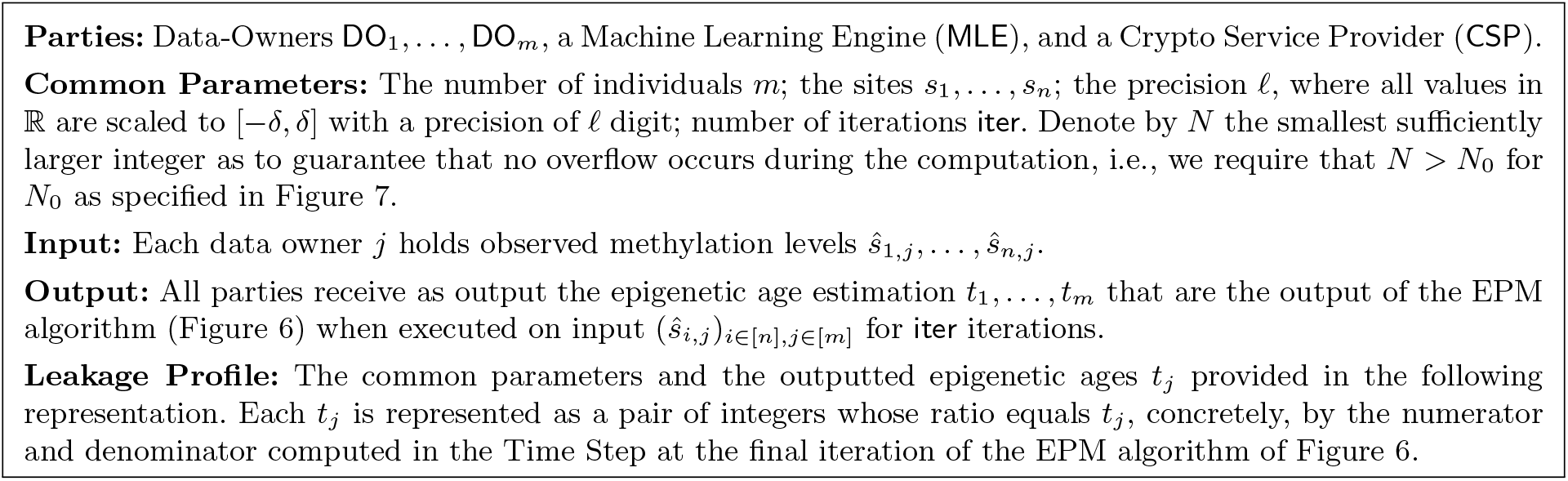
EPM Functionality, simplified to reveal the output

*Proof (of Theorem 1, Privacy, for the simplified functionality (Figure 9))*. Denote by *J* ⊆ [*m*] the set of data owners corrupted by the adversary. We consider three cases: the adversary controls also MLE or CSP or neither (controlling both servers is disallowed in the two-server model). Privacy in the third case follows immediately from any of the preceding two cases, because the MLE and CSP have no input and the output is public; we therefore focus on the first two cases.

*Case I – the adversary controls J and MLE*. We construct a ppt simulator Sim_1_ that receives the input of parties in *J* (MLE has no input), the outputted e-ages, and the leakage profile (public parameters and 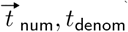), and produces a simulated-view for the adversary controlling *J* and MLE. The view includes the inputs of all corrupt parties, their random tape consisting of uniformly random values (of the required length) sampled by Sim_1_, and a simulated view of the messages they receive throughout the protocol constructed by Sim_1_ constructed as follows:

– Sim_1_ honestly generates (*pk, sk*) ← KeyGen(1^*λ*^, 1^*N*^) and adds *pk* to the simulated view (in place of the message received from CSP during setup).
– For every honest DO_*j*_ (i.e., *j* ∈*/ J*), Sim_1_ generates *n* independent random encryptions of zero under *pk* and adds them to the view (in place of encrypted inputs received by MLE from the honest DO_*j*_’s during data upload).
– For every corrupt DO_*j*_ (i.e., *j* ∈ *J*), Sim_1_ encrypts computes ctxt_*j*_ ← Enc(*pk*, (*ŝ*_*i,j*_)_*i*∈[*n*]_) using the random coins specified in the random tape of DO_*j*_, and adds these ciphertexts to the view (in place of encrypted inputs received by MLE from the corrupt DO_*j*_’s during data upload).
– Sim_1_ adds to the view 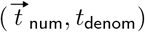 (in place of the e-ages published by CSP in the output post-processing phase).

We prove that the joint distribution of the real view together with the output of the protocol is computationally indistinguishable from the joint distribution of the simulated view together with the output of the functionality. This is proven via the following hybrids.

*ℋ*_0_ is the joint distribution of the adversary’s view and protocol’s output in a real execution of the protocol.
*ℋ*_1_ is similar to *ℋ*_0_, except that, in the view, the random tape, public key, and messages received during the output phase are as in the simulated view. Observe that *ℋ*_0_ and *ℋ*_1_ are identically distributed (because the random tape is selected uniformly at random, the keys are generated using an honest execution of KeyGen and the simulated messages for the output are the output provided to the simulator.
*ℋ*_2_ is similar to *ℋ*_1_ except that we replace the output of the protocol by the output of the functionality 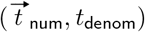, and also replace the output values published by the CSP upon the termination of the output post-processing phase by these same values 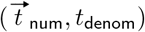 (as used the simulated view). Observe that *ℋ*_1_ and *ℋ*_2_ are identically distributed because the output of the protocol and the functionality are identically distributed (by the correctness of the protocol), which in turn implies that also the output values published by the CSP upon the termination of the output post-processing phase are identically distributed to the output of the functionality (here we rely on the fact that all parties receive the same output).
*ℋ*_3_ is the joint distribution of the simulated view (as computed by Sim_1_) and the output of the functionality. That is, *ℋ*_3_ is similar to *ℋ*_2_ except that the view is the simulated view, that is, the ciphertexts received from honest DO_*j*_’s are all replaced by encryptions of zero. By the semantic security of the encryption scheme, *ℋ*_2_ ≈_*c*_ *ℋ*_3_.

We conclude that *ℋ*_3_ ≈_*c*_ *ℋ*_0_. That is, the joint distribution of the simulated view and the output of the functionality is computationally indistinguishable from the joint distribution of the view of the adversary and the output of the protocol.

*Case II – the adversary controls J and CSP*. We construct a ppt simulator Sim_2_ that receives the public parameters, the input of parties in *J* (CSP has no input), the outputted e-ages and the leakage profiles, and produces a simulated-view for the adversary controlling *J* and CSP. The view includes the inputs of all corrupt parties, their random tape consisting of uniformly random values (of the required length) sampled by Sim_2_, and a simulated view of the messages they receive throughout the protocol constructed by Sim_2_ as follows.

– Sim_2_ executes (*pk, sk*) ← KeyGen(1^*λ*^, 1^*N*^) while using the randomness as set by Sim_2_ in the random tape of the CSP, and adds *pk* to the view (in place of the public key received by the corrupt DO_*j*_’s during setup).
– Sim_2_ computes 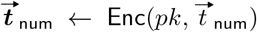 and ***t***_denom_ ← Enc(*pk, t*_denom_), and adds the ciphertexts 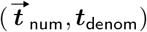 to the view (in place of the ciphertexts published by the MLE upon the termination of the secure computing phase).
– Sim_2_ adds to the view the cleartext values 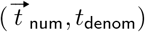 (in place of the output values published by the CSP upon the termination of the output post processing phase).

We prove that the joint distribution of the real view together with the output of the protocol is computationally indistinguishable from the joint distribution of the simulated view together with the output of the functionality. This is proven via the following hybrids.

*ℋ*_0_ is the joint distribution of the adversary’s view and protocol’s output in a real execution of the protocol.
*ℋ*_1_ is similar to *ℋ*_0_, except that, in the view, the random tape, public key. Observe that *H*_0_ and *H*_1_ are identically distributed (because the random tape is selected uniformly at random, the keys are generated using an honest execution of KeyGen and are consistent with the random tape of the CSP as set by Sim_2_, and the simulated messages for the output are the output provided to the simulator.
*ℋ*_2_ is the joint distribution of the simulated view (as computed by Sim_2_) and the output of the functionality. That is *ℋ*_2_ is similar to *ℋ*_1_ except that we replace the output of the protocol by the output of the functionality 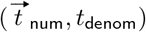, and also: replace the cleartext values published by the CSP upon the termination of the output post-processing phase by these same values 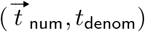, and replace the ciphertexts published by the MLE upon the termination of the secure computing phase by the encryption of these values using *pk*, i.e., by 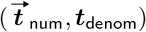 defined by 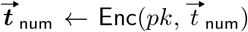 and ***t***_denom_ ← Enc(*pk, t*_denom_). Observe that *ℋ*_1_ and *ℋ*_2_ are identically distributed because the output of the protocol and the functionality are identically distributed (by the correctness of the protocol), which in turn implies that also the output values published by the CSP upon the termination of the output post-processing phase are identically distributed to the output of the functionality (here we rely on the fact that all parties receive the same output).

We conclude that *ℋ*_2_ ≈_*c*_ *ℋ*_0_. That is, that the joint distribution of the simulated view and the output of the functionality is computationally indistinguishable from the joint distribution of the view of the adversary and the output of the protocol.

*Proof (of Theorem 1, Privacy)*. We next sketch how we extend the above proof to show that our full protocol (Section 3.2) securely realizes the EPM functionality (Figure 1).

We first analyze the extension of the protocol for hiding the outputted numerators. The proof of privacy extends for the case of hiding 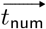 (see Section 3.2) by letting the simulator generate uniformly random values in place of the masked values for the honest data owners – resulting in an identical distribution (since the adversary has no information of what mask is generated by honest data owners). The formal proof is via extending the hybrid argument accordingly.

We next analyze the protocol extension for hiding both the numerators and the denominator. Extending the proof of privacy for the case of hiding also *t*_denom_ is done by considering two cases: either the adversary controls MLE or CSP (recall that it cannot control both). In case the adversary controls MLE (and any number of the data owners), the simulator generates a fresh encryption of zero in place of the ciphertext 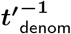 received by MLE from the CSP–relying on the semantic security of the FHE scheme–,and uses the output values *t*_*j*_ to simulate the messages received by the corrupt data owners DO_*j*_ inthe output post processing phase (recall that the simulator is provided the input and output of corrupt parties). In case the adversary controls CSP (and any number of the data owners), the simulator uses a uniformly random value in place of the 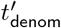 computed by MLE in a real execution of the protocol – resulting in identical distributions.

### A.4 Proof of Theorem 2 – Complexity

*Proof (of Theorem 2, Size)*.

Homomorphically computing *ψ*_*k*_ takes *m* + 1 homomomorphic additions and at most 2 homomomorphic multiplications.

Homomorphically computing *U* will therefore take *O*(*m ·* (*m* + 3)) = *O*(*m*^2^ + 3*m*) homomomorphic multiplications and addition operations.

Homomorphically computing *ϕ*_*k*_ takes 2*m* homomomorphic additions, 1 homomomorphic subtractraction, and 1 homomomorphic multiplication.

So, homomorphically computing *L* will take *O*(*m ·* (2*m* + 2) = *O*(2*m*^2^ + 2*m*) homomomorphic multiplications and addition operations.

Homomorphically computing 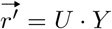 and 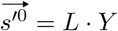 is done as follows to utilize the sparsity of *U* and *L*: Multiplying each row of *U* or *L* by *Y* requires computing only *m* homomorphic multiplication and addition operations (because the row has only *m* non-zero entries). So, doing so for all *n* row takes *nm* homomorphic operations.

Specifically, computing *r*^*′*^ will take in total *O*(*m*^2^ + 3*m* + 2*mn*) = *O*(*m*^2^ + *m*(3 + 2*n*)) and computing 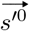 will take *O*(2*m*^2^ + 2*m* + 2*mn*) = *O*(2*m*^2^ + 2*m*(1 + *n*))

A summary of the site step total time complexity will be: *O*(3*m*^2^ + *m*(5 + 4*n*))

Computing *Λ*^−1^ takes *O*(2*m* + 1) homomorphic additions and multiplications.

Computing *t*_num_ takes *O*(*n*) homomorphic additions and multiplications.

Computing *t*_denom_ takes *O*(*n*) homomorphic additions and multiplications.

A summary of the time step total complexity will be: *O*(2*n* + 2*m* + 1) homomorphic operations.

Adding the complexity of both steps results in the following for a single iteration of the algorithm: *O*(3*m*^2^ + *m*(7 + 4*n*) + 2*n* + 1)

*Proof (of Theorem 2, Depth)*. For each variable *x* in our protocol, we denote the ×-depth of computting its value in iteration *k* by d(*x*; *k*). We show by induction that in each iteration 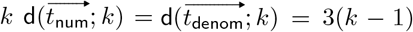. The base case is the first iteration. In this iteration, *t* consists of random values generated by the MLE and processed in cleartext. So, 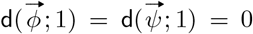, implying that 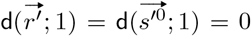 (where we assume that the underlying FHE supports plaintext vs. ciphertext multiplication being processed using homomorphic additions only as to incur no added ×-depth), and also d(*Λ*^−1^; 1) = 0; so, d(*t*_num_; 1) = d(*t*_denom_; 1) = 0.

Consider next the *k*th iteration for *k >* 1. The variables 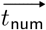 and *t*_denom_ entering this iteration are in encrypted form. Denote 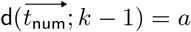 and d(*t*_denom_; *k* − 1) = *b*. So,

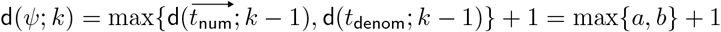

and

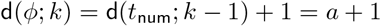

implying that

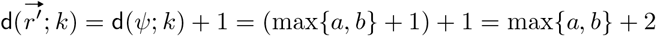

and

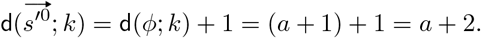

Likewise,

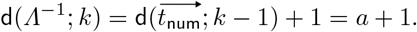

So,

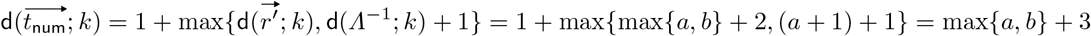

and

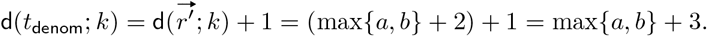

By the induction hypothesis *a* = *b* = 3(*k* − 1). Assigning these value for *a, b*, we get that 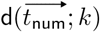 and d(*t*_denom_; *k*) are equal to max{*a, b*} + 3 = 3(*k* − 1) + 3 = 3*k*, which proves the desired.

### A.5 Proof of Maximum Plaintext Modulus Value 7

*Proof*. We define *N*_0_ as the largest integer value produced by the secure computation. We claim this to be one of the time step outputs (*t*_num_ or *t*_denom_), as the calculation in this step incorporates all calculations performed in the site step. We proceed to define the maximum possible values: 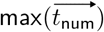 and max(*t*_denom_). In all calculations we must take into account that the age values represent the epigenetic state, and therefore may also be negative.

We denote *τ* as the maximal age value, *σ* as the maximal methylation value and (*υ ar*; *k*) as the value of the variable *var* in iteration *k*

– 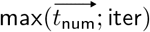: As depicted in 7, we first an upper bound for the maximum value of 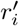 and 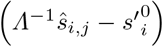
  - We begin by defining the maximal value in the vector 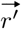. By inspecting the formula we conclude that the maximum value can be calculated as: *m ·* (2*m*(*τ* ; iter)) *· σ ·* (*t*_denom_; iter − 1)
  - Next we wish to maximize *Λ*^−1^*ŝ*_*i,j*_. We find the maximum value here to be: (*m*(*τ* ; iter))^2^ *· σ*
  - Due to the negative sign before 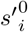, we wish to express the minimal possible value in the vector 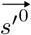. We define this value as: *m ·* (−2*m ·* (*τ* ; iter)^2^) *· σ* # «
  - Concluding from the above, we can define that 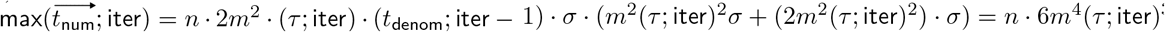
– 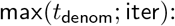 As depicted in 7, we wish to maximize the value 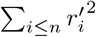 In order to achieve this, we can use the definition of 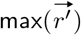 from the previous step. We then conclude that: 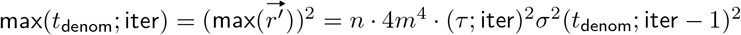

We now need to determine which of the values is larger. In the first iteration, we define (*t*_denom_; iter − 1) = 1, therefore, for this step it is clear that the value of 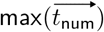 is the larger one as: *n ·* 6*m*^4^ *·*(*τ* ; iter)^3^*σ*^2^ *·* 1 *> n ·* 4*m*^4^ *·* (*τ* ; iter)^2^*σ*^2^ *·* 1.

For the next iteration, we set 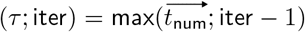 as this is now the largest age value. We then proceed to evaluate the values of 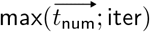 ans max(*t*_denom_; iter) as following:

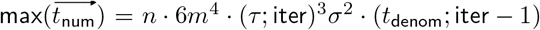

max(*t*_denom_; iter) = *n ·* 4*m*^4^ *·* (*τ* ; iter)^2^*σ*^2^ *·* (*t*_denom_; iter − 1)^2^

We now need to determine whether 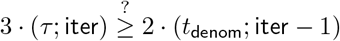

The above can be written as: 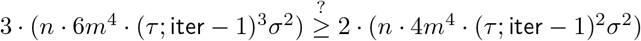

From the above it is clear that 9 *·* (*τ* ; iter− 1) *>* 4

And therefore we conclude that 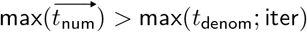 □

### A.6 Homomorphic Encryption and Chinese Remainder Theorem

As mentioned in Section 3, performing arithmetic operation on rational numbers represented as pairs of “numerator, denominator” causes (plaintext) values occurring in the algorithm to rapidly increase, beyond the point they can be accurately represented. To address this issue, we employ the Chinese Remainder Theorem (CRT), following [4]. Essentially this is done as follows (see details in [4]).

*𝒫* = (*p*_1_, *p*_2_, …, *p*_*n*_) be a sequence of co-prime integers and *X* ∈ {0, …, Π _*i*_ *p*_*i*_ − 1}, then by CRT *X* can be uniquely represented as (*x*_1_, *x*_2_, …, *x*_*n*_), where:

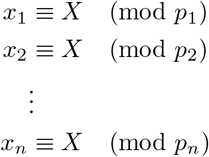

Moreover, the inverse process is efficient and given by:

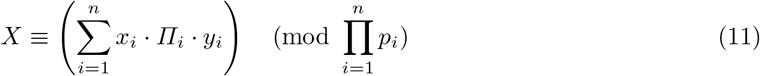

where: 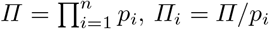, and *y*_*i*_ *·* Π_*i*_ ≡ 1 (mod *p*_*i*_).

Moreover, using the representation defined above (also known as the Residue Number System representation (RNS)), has additional advantages such as amneability to parallelization of computation.For example, let 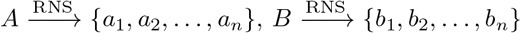 then the result of some operation *C* is defined as follows:

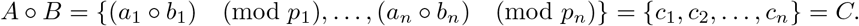

where ∘ is some arithmetic operation.

In the context of an FHE scheme (KeyGen, Enc, Dec, Eval) the CRT representation can be employed as follows.

– Key generation is done by generating a key pair (*pk*_*i*_, *sk*_*i*_) ← KeyGen(1^*λ*^, *p*_*i*_) for each *i* and outputting the key pair 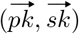 where 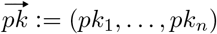 and 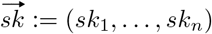.
– To encrypt a message *x* ∈ {0, …, Π *p*_*i*_ − 1}, compute *c*_*i*_ ← Enc(*pk*_*i*_, *x* mod *p*_*i*_), for all *i*, and output 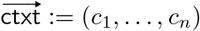.
– To decrypt (*c*_1_, …, *c*_*n*_) compute *m*^*′*^ ← Dec(*sk*_*i*_, *c*_*i*_), for all *i*, and output 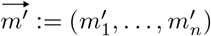.
– Homomorphic evaluation of a function *C* over *k* inputs given ciphertexts 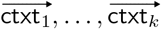(where each 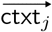 is a CRT vector 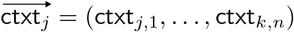), is done by computing 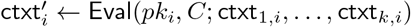, for each *i*, and outputting 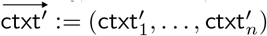.

The correctness of decryption and homomorphic evaluation and the semantic security all follow immediately from these properties for the underlying scheme.

## Footnotes

1 We refer here to our input-hiding protocol (cf. Section 3.1), which addresses the same privacy concern as in [16]. The stated complexity is of the secure computing phase in both ours and [16]’s protocols, while disregarding their pre- and post-processing phases because they are simple and comparable (consisting of key setup, input upload, and output post-processing); see the full complexity analysis in Theorem 2.

2 In further detail, we present three variants of our protocol. In two of those variants the role of the 2nd server is exactly that of a KMS. In the third variant, this 2nd server executes one additional computation: computing the modular inverse of a single (masked) output value.

3 All measurements are for a known and identical set of genome sites.

4 See [17] Chapter 3.2 for the standard notion of computational indistinguishable.

5 For the simplicity of representation, we assume that each DO holds a single methylation value for a single individual; in practice, a data owner may hold methylation values for an arbitrary number of individuals.

6 Alternatively, the CSP can provide the secret decryption key to any party (other than the MLE) who is authorized to decrypt and obtain the output. In this case, standard techniques must be employed to enroll and authenticate users, using proper credentials, to enforce the permissions policy.

## References

1. BFV context and key setup. https://pyfhel.readthedocs.io/en/latest/_autoexamples/Demo_2_Integer_BFV.html#sphx-glr-autoexamples-demo-2-integer-bfv-py. Accessed: 2024-01-17.

2. A. Akavia, B. Galili, H. Shaul, M. Weiss, and Z. Yakhini. Efficient privacy-preserving viral strain classification via k-mer signatures and FHE. In IEEE 36th Computer Security Foundations Symposium (CSF), pages 178–193. IEEE Computer Society, 2023.

3. A. Akavia, B. Galili, H. Shaul, M. Weiss, and Z. Yakhini. Privacy preserving feature selection for sparse linear regression. Proc. Priv. Enhancing Technol., 2024(1):300–313, 2024.

4. A. Akavia, H. Shaul, M. Weiss, and Z. Yakhini. Linear-regression on packed encrypted data in the two-server model. In Proceedings of the 7th ACM Workshop on Encrypted Computing & Applied Homomorphic Cryptography, WAHC’19, page 21–32, New York, NY, USA, 2019. Association for Computing Machinery.

5. A. Akavia, H. Shaul, M. Weiss, and Z. Yakhini. Linear-regression on packed encrypted data in the two-server model. In M. Brenner, T. Lepoint, and K. Rohloff, editors, Proceedings of the 7th ACM Workshop on Encrypted Computing & Applied Homomorphic Cryptography, WAHC@CCS 2019, London, UK, November 11-15, 2019, pages 21–32. ACM, 2019.

6. E. Al-Radadi and P. Siy. Rns sign detector based on chinese remainder theorem ii (crt ii). Computers & Mathematics with Applications, 46(10-11):1559–1570, 2003.

7. M. Blatt, A. Gusev, Y. Polyakov, and S. Goldwasser. Secure large-scale genome-wide association studies using homomorphic encryption. Proceedings of the National Academy of Sciences, 117(21):11608–11613, 2020.

8. C. Bonte, E. Makri, A. Ardeshirdavani, J. Simm, Y. Moreau, and F. Vercauteren. Towards practical privacy-preserving genome-wide association study. BMC bioinformatics, 19(1):1–12, 2018.

9. Z. Brakerski. Fully homomorphic encryption without modulus switching from classical gapsvp. In Annual Cryptology Conference, pages 868–886. Springer, 2012.

10. S. Carpov, N. Gama, M. Georgieva, and D. Jetchev. GenoPPML – a framework for genomic privacy-preserving machine learning. In 2022 IEEE 15th International Conference on Cloud Computing (CLOUD), pages 532–542, 2022.

11. C. clarity in privacy. The genetic information privacy act, 2021.

12. C. Dong, J. Weng, J.-N. Liu, A. Yang, L. Zhiquan, Y. Yang, and J. Ma. Maliciously secure and efficient large-scale genome-wide association study with multi-party computation. IEEE Transactions on Dependable and Secure Computing, 2022.

13. J. Fan and F. Vercauteren. Somewhat practical fully homomorphic encryption. Cryptology ePrint Archive, 2012.

14. P. Fouque, J. Stern, and J. Wackers. Cryptocomputing with rationals. In FC’02, pages 136–146, 2002.

15. C. Gentry. A fully homomorphic encryption scheme. PhD thesis, Stanford University, 2009. crypto.stanford.edu/craig.

16. M. Goldenberg, S. Snir, and A. Akavia. Private epigenetic pacemaker detector using homomorphic encryption. In International Symposium on Bioinformatics Research and Applications, pages 52–61. Springer, 2022.

17. O. Goldreich. The Foundations of Cryptography - Volume 1: Basic Tools. Cambridge University Press, 2004.

18. S. Halevi and V. Shoup. Algorithms in HElib. In Annual Cryptology Conference, pages 554–571. Springer, 2014.

19. C. R. Harris, K. J. Millman, S. J. van der Walt, R. Gommers, P. Virtanen, D. Cournapeau, E. Wieser, J. Taylor, S. Berg, N. J. Smith, R. Kern, M. Picus, S. Hoyer, M. H. van Kerkwijk, M. Brett, A. Haldane, J. F. del Río, M. Wiebe, P. Peterson, P. Gérard-Marchant, K. Sheppard, T. Reddy, W. Weckesser, H. Abbasi, C. Gohlke, and T. E. Oliphant. Array programming with NumPy. Nature, 585(7825):357–362, Sept. 2020.

20. S. Hong, J. H. Park, W. Cho, H. Choe, and J. H. Cheon. Secure tumor classification by shallow neural network using homomorphic encryption. BMC genomics, 23(1):1–19, 2022.

21. S. Horvath. Dna methylation age of human tissues and cell types. Genome Biology, 14(10):3156, 2013.

22. A. Ibarrondo and A. Viand. Pyfhel: Python for homomorphic encryption libraries. In Proceedings of the 9th on Workshop on Encrypted Computing & Applied Homomorphic Cryptography, pages 11–16, 2021.

23. A. E. Jaffe, Y. Gao, A. Deep-Soboslay, R. Tao, T. M. Hyde, D. R. Weinberger, and J. E. Kleinman. Mapping dna methylation across development, genotype and schizophrenia in the human frontal cortex. Nature neuroscience, 19(1):40–47, 2016.

24. W.-J. Lu, Y. Yamada, and J. Sakuma. Privacy-preserving genome-wide association studies on cloud environment using fully homomorphic encryption. In BMC medical informatics and decision making, volume 15, pages 1–8. Springer, 2015.

25. A. Meurer, C. P. Smith, M. Paprocki, O. Čertík, S. B. Kirpichev, M. Rocklin, A. Kumar, S. Ivanov, J. K. Moore, S. Singh, T. Rathnayake, S. Vig, B. E. Granger, R. P. Muller, F. Bonazzi, H. Gupta, S. Vats, F. Johansson, F. Pedregosa, M. J. Curry, A. R. Terrel, v. Roučka, A. Saboo, I. Fernando, S. Kulal, R. Cimrman, and A. Scopatz. Sympy: symbolic computing in python. PeerJ Computer Science, 3:e103, Jan. 2017.

26. S. of California Department of Justice. California consumer privacy act (CCPA), 2018.

27. J. of the European Union. Regulation (EU) 2016/679 of the european parliament, 2016.

28. G. M. Pinho, J. G. Martin, C. Farrell, A. Haghani, J. A. Zoller, J. Zhang, S. Snir, M. Pellegrini, R. K. Wayne, D. T. Blumstein, et al. Hibernation slows epigenetic ageing in yellow-bellied marmots. Nature ecology & evolution, 6(4):418–426, 2022.

29. R. L. Rivest, L. Adleman, and M. L. Dertouzos. On data banks and privacy homomorphisms. Foundations of Secure Computation, Academia Press, pages 169–179, 1978.

30. C. F. Sagi Snir and M. Pellegrini. Human epigenetic ageing is logarithmic with time across the entire lifespan. Epigenetics, 14(9):912–926, 2019.

31. E. Shiriaev, N. Kucherov, M. Babenko, and A. Nazarov. Fast operation of determining the sign of a number in rns using the akushsky core function. Computation, 11(7):124, 2023.

32. S. Simmons and B. Berger. Realizing privacy preserving genome-wide association studies. Bioinformatics, 32(9):1293–1300, 2016.

33. S. Snir. Epigenetic pacemaker: closed form algebraic solutions. BMC genomics, 21(2):1–11, 2020.

34. S. Snir, C. Farrell, and M. Pellegrini. Human epigenetic ageing is logarithmic with time across the entire lifespan. Epigenetics, 14(9):912–926, 2019.

35. S. Snir and M. Pellegrini. An epigenetic pacemaker is detected via a fast conditional expectation maximization algorithm. Epigenomics, 10(6):695–706, 2018.

36. S. Snir, B. M. vonHoldt, and M. Pellegrini. A statistical framework to identify deviation from time linearity in epigenetic aging. PLoS computational biology, 12(11):e1005183, 2016.

37. S. Snir, Y. I. Wolf, and E. V. Koonin. Universal pacemaker of genome evolution. PLoS computational biology, 8(11):e1002785, 2012.

38. S. Snir, Y. I. Wolf, and E. V. Koonin. Universal pacemaker of genome evolution in animals and fungi and variation of evolutionary rates in diverse organisms. Genome biology and evolution, 6(6):1268–1278, 2014.

39. P. S. Wang, M. J. T. Guy, and J. H. Davenport. P-adic reconstruction of rational numbers. ACM SIGSAM Bulletin, 16(2):2–3, 1982.

40. Y. I. Wolf, S. Snir, and E. V. Koonin. Stability along with extreme variability in core genome evolution. Genome biology and evolution, 5(7):1393–1402, 2013.

41. J. Zhou, B. Lei, and H. Lang. Homomorphic multi-label classification of virus strains. In 2022 IEEE International Symposium on Software Reliability Engineering Workshops (ISSREW), pages 289–294. IEEE, 2022.

